# Homophilic and heterophilic cadherin bond rupture forces in homo- or hetero-cellular systems measured by AFM based SCFS

**DOI:** 10.1101/2020.02.11.943597

**Authors:** Prem Kumar Viji Babu, Ursula Mirastschijski, Gazanfer Belge, Manfred Radmacher

## Abstract

Cadherins enable intercellular adherens junctions to withstand tensile forces in tissues, e.g. generated by intracellular actomyosin contraction. Single molecule force spectroscopy experiments in *in-vitro* experiments can reveal the cadherin-cadherin extracellular region binding dynamics such as bond formation and strength. However, characterization of cadherin homophilic and heterophilic binding in their native conformational and functional state in living cells has rarely been done. Here, we used Atomic Force Microscopy (AFM) based Single cell force Spectroscopy (SCFS) to measure rupture forces of homophilic and heterophilic bond formation of N-, OB- and E-cadherins in living fibroblast and epithelial cells in homo- and hetero-cellular arrangements, i.e. between same type of cells and between cells of different type. In addition, we used indirect immunofluorescence labelling to study and correlate the expression of these cadherins in intercellular adherens junctions. We showed that N/N and E/E cadherin homophilic bindings are stronger than N/OB, E/N and E/OB heterophilic bindings. Disassembly of intracellular actin filaments reduces the cadherin bond rupture forces suggesting a contribution of actin filaments in cadherin extracellular binding. Inactivation of myosin did not affect the cadherin rupture force in both homo- and hetero-cellular arrangements. Whereas, myosin inactivation particularly strengthened the N/OB heterophilic bond and reinforced the other cadherins homophilic bonds.

## Introduction

Cell adhesion to neighbouring cells or the extracellular matrix (ECM) environment is a very important process in regulating crucial biological activities such as embryonic development, tissue assembly and dynamics, wound healing and cancer metastasis. Generally, cells communicate with other cells through adherens, gap or mechanosensitive junctions (1). Cadherins from adherens junctions are a class of calcium dependent cell adhesion molecules (CAMs) which comprise three different domains: (i) an intracellular or cytoplasmic domain which binds to the actin cytoskeleton through adaptor proteins such as α-catenin, β-catenin and p120 catenin, (ii) a transmembrane domain and (iii) an extracellular domain. The extracellular domain consists of five extracellular cadherin (EC) repeats. A dimer of EC1-EC5 of one cell interacts with the corresponding cadherin dimer of a neighbouring cell through homophilic or heterophilic interaction (2, 3).

Several assays have been developed to investigate cell-cell interactions in the last two decades, such as dual micropipette assay (4), flipping assay (5), FRET (6) and AFM based SCFS (7, 8). Comparing all assays, the AFM based SCFS assay provides a wide range of forces (10 pN to 10^6^ pN) (9) and a controlled force application (loading rate) on the cell-cell adhesion cadherin bond by retracting the AFM cantilever at a well-defined velocity (10). In SCFS, cell adhesion force measurements are performed in near physiological conditions. Being a multifunctional toolbox in nanobiotechnology (11), AFM provides a functionalized cantilever to pick up a live cell guided by optical microscopy. It allows probing the rupture force between cadherin molecules present in two cells, by separating the two cells. The rupture force can be quantified and reveals differences in the specific type of cadherins secreted by different cell types.

According to the presence or absence of the HAV (His-Ala-Val) cell recognition sequence in the EC1 domain, classical cadherins are classified into type I (E-, N- and others) and type II cadherins (OB- and others) (2,3). The most commonly expressed cadherin found in fibroblasts is N-cadherin (cad-2) (12). Primary rat fibroblasts differentiate into myofibroblasts *in vitro* using transforming growth factor- β1 (TGF-β1). TGF- β1 induces the expression of alpha-smooth muscle actin (α-sma), an increased expression of OB-cadherins (cad-11) and a decreased expression of N-cadherin (13). This TGF-β1 induced cadherin switch from N-cadherin to OB-cadherin increases the intercellular adhesion strength between myofibroblasts by strengthening individual OB-cadherin bonds. Single molecule force spectroscopy (SMFS) measurements on OB- and N-cadherins showed that the rupture force between OB-cadherins homophilic interaction is larger than between N-cadherins (14). A biochemical analysis of N- and OB-cadherins expression in human dermal fibroblast and Dupuytren’s myofibroblast shows increased OB-cadherin and decreased N-cadherin expression in myofibroblasts compared to dermal fibroblasts (1). The E (epithelial)-cadherin (cad-1) is the dominant cadherin expressed in most epithelial cell lines like MDCK (Madine-Darby Canine Kidney) cells (15). The more motile, trypsin sensitive subpopulation of MDCK cells shows a low level of N-cadherin expression (16).

Hetero-cellular interactions between different cell types occur in tissue and organ morphogenesis. Involvement of specific cadherins in these interactions plays a pivotal role in cancer cell metastasis (17) whereas heterophilic interactions between cell specific cadherins mediate cancer cell invasion (18). Direct interactions between fibroblast and epithelial cells may play an important role in the epithelial to mesenchymal transition (EMT) process (19). Hetero-cellular interactions between normal fibroblasts and gastric cancer cells induce E-cadherin loss and increase metastasis in gastric cancer via EMT (20). The investigation of hetero-cellular interactions between fibroblast and epithelial cells using biophysical techniques such as SCFS will help to better understand the role of classical cadherin interactions both in EMT and Mesenchymal to Epithelial cell transition (MET) processes.

Actin filaments associated with myosin are the major contractile component responsible for intracellular force generation. Generally, these forces are generated by the myosin assembly and motility on the actin filaments. Myosin light chain is phosphorylated by the myosin light chain kinase (MLCK) and this activates the myosin cross linking to the actin filaments with actomyosin contractile force generation. Intracellular forces are then transmitted to the neighbouring cells and to the extracellular environment through cadherins and integrins, respectively that are connected to actin filaments. Disassembling actin filament rich stress fibres by treating fibroblasts with Cytochalasin D results in decreased cell stiffness (21). Addition of Cytochalasin D reduces the cadherin mediated binding forces between myofibroblasts, as measured by SCFS, and shows that cadherins are linked structurally and possibly functionally to the intracellular actin network (14). Inactivating myosin-II activity by treating fibroblasts with ML-7 inhibits the MLCK, which further prevents myosin mediated actomyosin contractility which results in actin cytoskeleton softening and thus decreased cell stiffness (22).

In the present study, we have studied the expression of N- and OB-cadherins in three types of fibroblasts extracted from the same patient with Dupuytren’s disease using fluorescence microscopy: (1) normal fibroblasts-NFs from normal healthy skin, (2) scar fibroblasts-SFs from cutaneous scar tissue and (3) Dupuytren’s myofibroblast-DFs from the nodules of the palmar fascial strands. Using AFM-SCFS, we measured the rupture forces between fibroblasts grown in a confluent monolayer and fibroblasts attached to the AFM cantilever (NF-NF, SF-SF and DF-DF). Loading rate dependent rupture force measurements showed that NF and SF exhibit larger rupture forces than DFs. These results correlated with the cadherin types present in the adherens junctions of respective fibroblast types. Hetero-cellular interaction forces were also measured between fibroblasts grown in monolayers and epithelial cells attached to the cantilever. Regarding the epithelial cell, we used epithelial cell line called MDCK cells to study the hetero-cellular interactions between MDCK and fibroblasts mediated by cadherins expression and binding dynamics. Immunofluorescence studies of MDCK and fibroblast co-cultures showed the presence of N-cadherins at the fibroblast-MDCK junction and E-cadherin loss in MDCK. Cytochalasin D treatment decreases the interaction forces in both homo-cellular and hetero-cellular interactions. In ML-7 treatment, no change in interaction forces observed in homo-cellular and hetero-cellular interactions except for DF-DF interaction. Contrarily, there is an increase in DF-DF rupture forces after ML-7 treatment and reveals that OB- and N-cadherin heterophilic bond strengthens the cell-cell interaction when there is no intracellular contractile force.

## Results

### N/OB heterophilic binding is weaker than N- and OB-homophilic binding

Investigation of cell-cell interactions using AFM becomes more possible using a simple cell force spectroscopic setup. AFM based SCFS setup is explained with the simple schematics shown in Fig 1A. A tipless cantilever, functionalized with concanavalin A (conA), was placed on a cell, which makes initial adhesion to the substrate and is appropriately round in shape. The cantilever was approached towards that cell until a certain loading force has been reached. After a dwell time of 5 sec the cell has adhered sufficiently and stays attached to the cantilever when the cantilever is retracted from the support as shown in Fig. 1B. The force curve obtained during cell capture is shown in Fig. 1C. After a recovery time of 10 minutes, the cantilever with the attached cell was approached towards and retracted from another cell attached to the Petri dish. Cell-cell interactions and rupture forces between cells were probed. Fig. 1D shows a cell-cell (NF-NF) interaction force curve. The force curve contains approach (red arrows) and retract curve (blue arrows). The cell capturing and cell-cell inter-action events are visible in the retract curve. In case of cell-cell interactions, two distinct features can be seen in the retract curve: rupture (continuous line arrows) and tether events (discontinuous line arrows). The adhesion molecules that are well anchored to the intracellular actin filaments interact with their counterparts on the other cell, and the breakage of the adhesion molecules mechanical bonds can be seen as a rupture event. This rupture event can be due to a single bond breakage or to multiple bond breakages. The rupture force was calculated from the height of the rupture event. When adhesion molecules are not anchored to actin filaments membrane, tethers can be pulled over large distances, which eventually will also break (tether events). The rupture and tether events observed during cell capture were due to the interaction and bond breakage of either specific adhesion molecules or other non-specific interactions, which were not characterized here.

**Figure 1.**
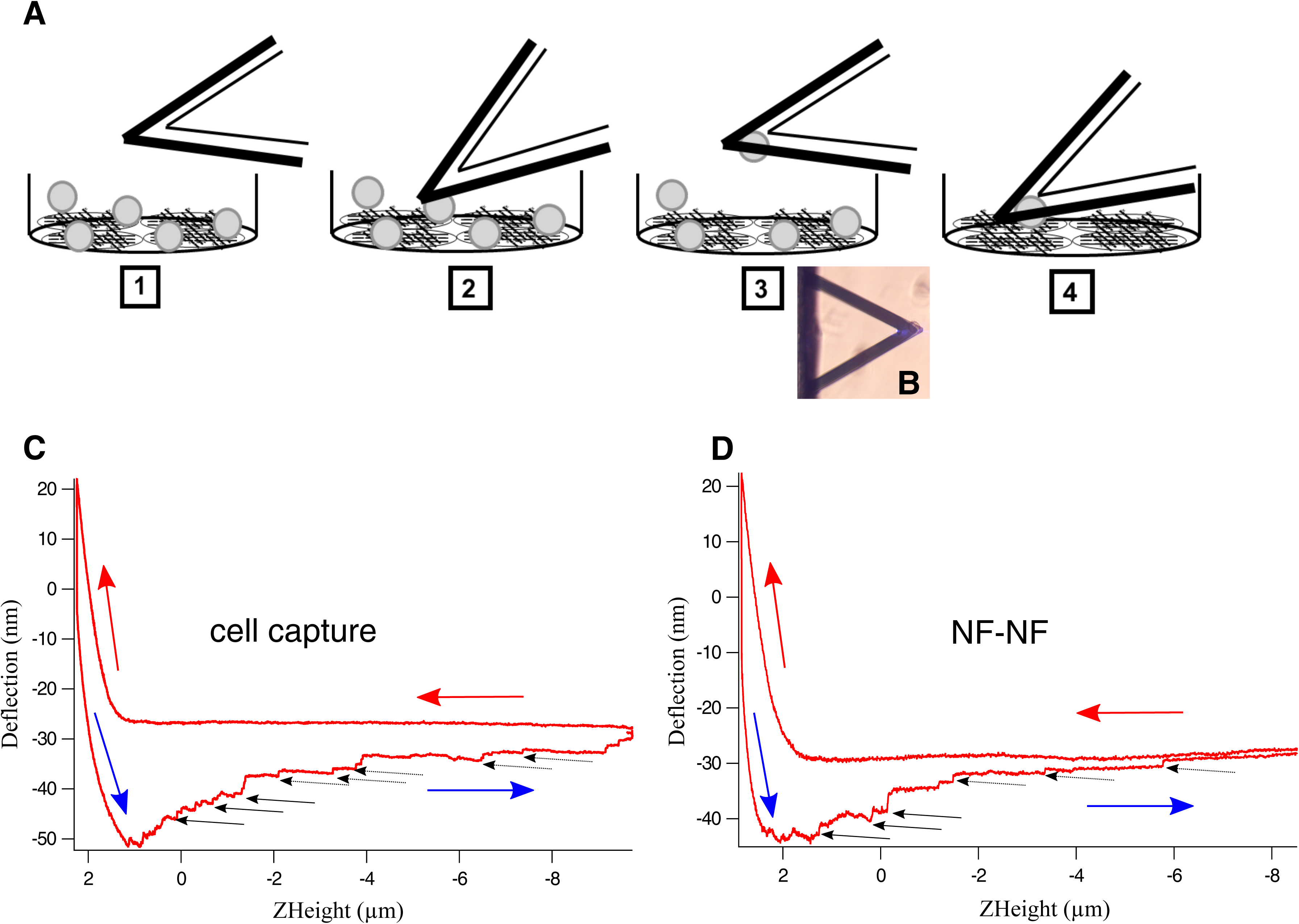
Schematic representation AFM based SCFS experimental setup. (A) This cartoon represents the capturing of cell by a tipless cantilever in a stepwise manner. 1-The conA functionalized tipless cantilever and a cell with round morphology is chosen with the aid of optical microscopy. 2-The cantilever is approached towards the cell at certain velocity (5 μm/sec) and contact force (3.5 nN). 3-Given the contact time of 5 s, the cell attached cantilever is retracted with the same velcotiy (5 μm/sec). (B) Optical image shows the cell attached cantilever. (C) Force curve recorded during cell pick up was shown and retract curve (blue arrow) contains rupture (continuous line black arrow) due to unspecific binding and tether (discontinuous line black arrow) events. 4- After a recovery time of 10 min, the interaction between cell attached to the cantilever and cell grown as monolayer was conducted. (D) Cell-cell interaction force curve shows rupture events that corresponded to the extracellular cadherin-cadherin bond breakage. Here, multiple rupture events were recorded.

Here we determined rupture forces between three types of fibroblasts isolated from primary human cells using SCFS and assessed the specific cadherins at the interaction site using fluorescence microscopy. The cell-cell rupture force was measured using an approach and retraction velocity of 3 μm/sec, a maximum loading force of 3 nN and the contact time of 2 s. The histogram plot of measured rupture forces versus the number of rupture events shows the force distribution for each fibroblast type (NF-NF Fig. 2A, SF-SF Fig. 2B and DF-DF Fig. 2C-red bar). NF-NF interaction showed a larger rupture force (51.91 pN) compared with SF-SF (45.21 pN) or DF-DF (35.71 pN) (See Table 1 which lists the corresponding 25, 50 and 75 percentile values). To verify that these rupture forces were due to the cadherin-cadherin bond breakages, the rupture events were recorded in the presence of EGTA (ethylene glycol tetraacetic acid, a calcium chelating agent) in the SCFS setup, effectively removing all free calcium from the extracellular space. Addition of EGTA completely inhibited the cadherin mediated cell-cell interaction with reduced numbers of rupture events (Fig. 2A, B and C, blue bar). Under normal conditions, force curves showed multiple rupture events due to interactions of multiple cadherins (Supplementary Fig. 1A), whereas in the absence of Ca^2+^, i.e. in the presence of EGTA, such rupture events were not seen in force curves (Supplementary Fig. 1B). To understand the cadherin-cadherin binding strength, we exerted varying force (loading) rates on the bonds by approaching and retracting the AFM cantilever at different velocities (3, 5, 7.5 and 10 μm/sec), which named “pulling rate” in force spectroscopy. For all three fibroblast types, the corresponding rupture forces showed a linear increase depending on the pulling rate applied (Fig. 2D). The median rupture force values for respective pulling rates for all three fibroblast types were listed in Table 1. NF-NF (Fig. 2D black square) and SF-SF (Fig. 2D red upper triangle) rupture forces were similar at all velocities except 3 μm/sec. In contrast, DF-DF (Fig. 2D blue lower triangle) attachments showed smaller rupture forces compared to NF-NF and SF-SF at all velocities.

**Table 1:** The median (bold values) rupture force (pN) values of cadherin mediated homocellular and heterocellular adherens junctions at each approach and retract velocities.

| Approach and<br>Retract velocity<br>($\mu\text{m}/\text{sec}$) | 3 | | | 5 | | | 7.5 | | | 10 | | |
| --- | --- | --- | --- | --- | --- | --- | --- | --- | --- | --- | --- | --- |
|  | 25th | **Median** | 75th | 25th | **Median** | 75th | 25th | **Median** | 75th | 25th | **Median** | 75th |
| Cell-cell interaction |  |  |  |  |  |  |  |  |  |  |  |  |
| NF-NF | 21.53 | **51.91** | 27.20 | 24.77 | **73.40** | 40.72 | 33.18 | **97.18** | 61.16 | 41.63 | **117.39** | 85.12 |
| SF-SF | 21.56 | **45.21** | 32.33 | 28.00 | **74.22** | 53.77 | 39.56 | **103.28** | 77.95 | 49.35 | **128.51** | 107.84 |
| DF-DF | 17.59 | **35.71** | 21.08 | 21.57 | **58.23** | 31.23 | 28.92 | **83.54** | 49.22 | 36.28 | **103.05** | 67.51 |
| MDCK-MDCK | 35.83 | **75.51** | 50.04 | 39.74 | **98.48** | 58.59 | 47.88 | **124.08** | 82.34 | 60.41 | **149.68** | 111.48 |
| NF- MDCK | 20.34 | **55.85** | 30.04 | 22.23 | **70.48** | 35.55 | 29.08 | **91.44** | 48.63 | 35.93 | **111.28** | 66.59 |
| SF- MDCK | 30.53 | **39.68** | 40.66 | 29.14 | **68.19** | 48.56 | 31.12 | **92.36** | 61.87 | 34.45 | **106.62** | 74.06 |
| DF- MDCK | 20.30 | **46.70** | 26.19 | 21.84 | **63.62** | 32.67 | 27.17 | **83.20** | 42.12 | 32.78 | **99.42** | 53.18 |

**Figure 2.**
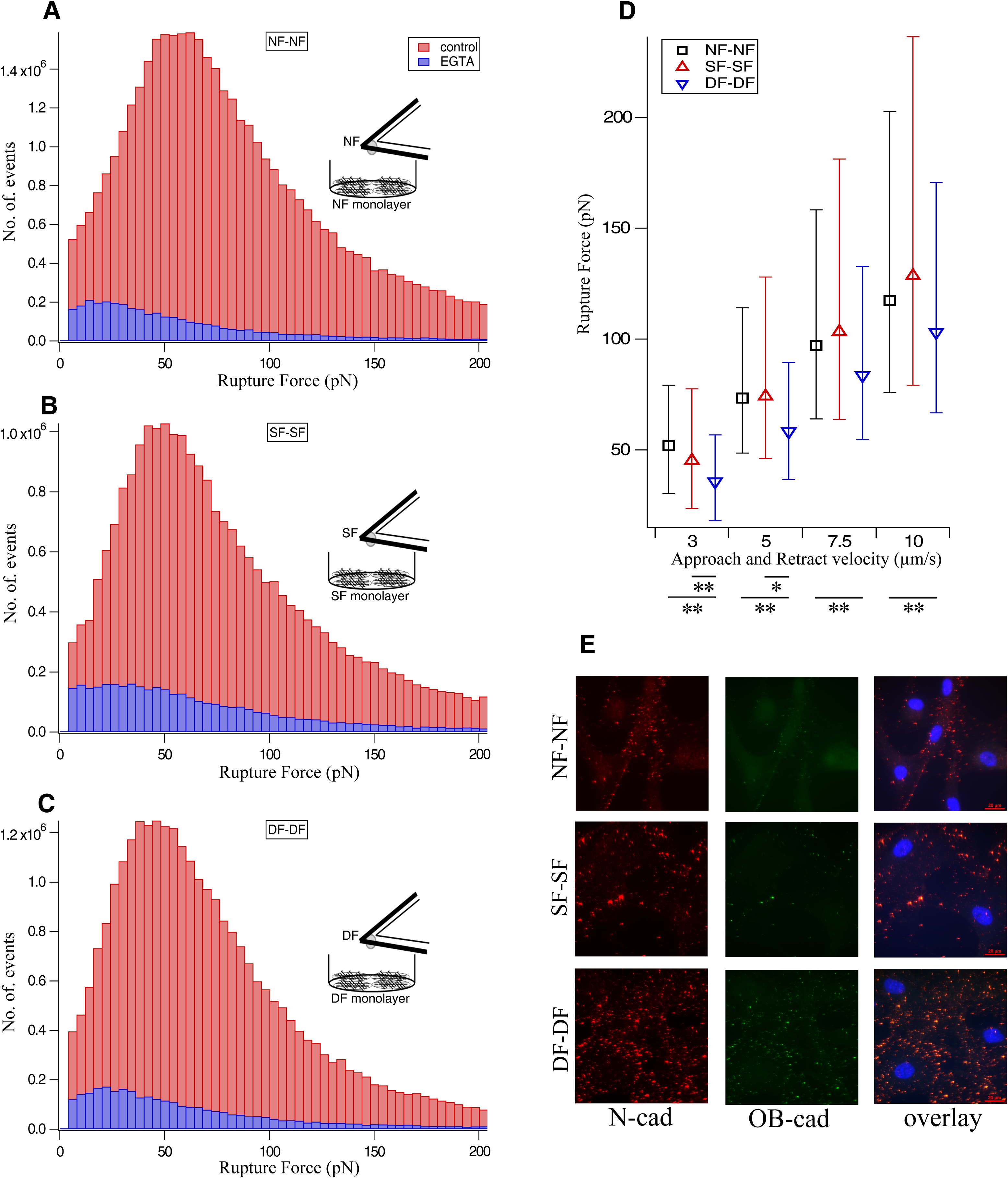
Fibroblast intercellular cadherin expression and rupture force measurement. Histogram shows the rupture force (red bar) recorded for (A) NF-NF, (B) SF-SF and (C) DF-DF interactions. Cadherin involvement in the rupture events (Supplementary Fig. 1A) was controlled by EGTA (blue bar) addition to the cell-cell interaction setup. This leads to the respective loss of rupture events (Supplementary Fig. 1B). (D) Increasing the approach and retract velocity of the cantilever linearly increases the cadherin rupture force for NF-NF (black square), SF-SF (red upper triangle) and DF-DF (blue lower triangle). NF and SF displays large rupture forces than DF at all velocities. (E) Dual immunofluorescence data shows N-cadherin expression (red fluorescence) in all fibroblasts adherens junctions. OB-cadherin expression (green fluorescence) was seen only in DF. The overlay (orange fluorescence) represents the heterophilic binding of N-cadherin and OB-cadherin which is encountered only in DF. Blue fluorescence indicates the nuclei. Scale bar 20 μm.

In order to assess the presence of specific cadherin types in cell-cell interaction sites, all three fibroblast types were immunostained for N- and OB- cadherin. Dual immunostaining for N- (red) and OB-(green) cadherins showed that NF and SF express exclusively N-cadherin whereas DF express both N- and OB- cadherin at the interaction site between cells (Fig. 2E). In the overlay (orange), heterophilic interactions between the N- and OB-cadherins are visible. Single immunostaining for N-cadherin (red) showed that all three fibroblast types express N-cadherin at the interaction site (Supplementary Fig. 2A). Similarly, single staining for OB-cadherin (red) in all fibroblasts showed that only DF-DF express OB-cadherin with absent OB-cadherin expression found at the NF-NF and SF-SF interaction site (Supplementary Fig. 2B). Controls with no primary antibody for N- (Supplementary Fig. 2C) and OB-cadherins (Supplementary Fig. 2D) shows no fluorescence that proves no unwanted or unspecific binding of fluorescently tagged secondary antibodies, thus showing the specificity of the secondary antibodies for the primary antibodies used here. This further confirms that the red and green fluorescence seen in Fig. 2E and Supplementary Fig. 2A&B are due to specific expression of N-cadherin in all three fibroblast types and OB-cadherin only in DF. This reveals homophilic N-cadherin binding in NF and SF and heterophilic N-cadherin/OB-cadherin binding in DF. Homophilic N-cadherin intercellular binding exhibited stronger interaction forces than N/OB-cadherin heterophilic binding when immunostaining results were compared to cadherin-cadherin bond rupture mechanical measurements.

### E-, N- and OB- cadherin at the fibroblast-epithelial hetero-cellular adherens junctions

The significance of studying hetero-cellular interactions may lead to sorting out different cell types by their expression and assembly of cell specific cadherins at the interaction site. The investigation of cadherin homophilic and heterophilic interactions may pave the way for a better understanding of cadherin mediated intracellular signalling. Heterophilic cadherin rupture forces were measured between epithelial cells and fibroblasts. A monolayer of fibroblasts was grown in a Petri dish and MDCK epithelial cells were attached to a tipless cantilever functionalized with conA. Fibroblast-MDCK interactions were studied by approaching a cantilever with attached MDCK cells towards the fibroblast cell monolayer at 3 μm/sec velocity with 3 nN maximum contact force and 2s contact time. In a similar fashion, MDCK-MDCK interactions were studied and the resulting median rupture force value was 75.51 pN. Regarding fibroblast-MDCK interaction, the median rupture force values were 55.85 pN for NF-MDCK, 39.68 pN for SF-MDCK and 46.70 pN for DF-MDCK (See Table 1 which lists the corresponding 25, 50 and 75 percentile values). In order to confirm the cadherin mediated rupture force, the force curves were recorded in the presence of EGTA. The histogram plot (Fig. 3A, B, C and D) showed a decrease in rupture events (blue bar) comparative to the rupture events (red bar) obtained without EGTA. The binding strength of the cadherins present in the membrane of these cell types was measured by approaching and retracting the cantilever with the attached MDCK cell at different velocities. All fibroblast-MDCK hetero-cellular interactions and also MDCK-MDCK binding (Fig. 3E) showed a linear increase in rupture force as a function of loading rate. The median rupture force values calculated for each pulling velocity for all three types of fibroblasts and MDCK or MDCK-MDCK interactions are listed in Table 1. Comparing the rupture forces, NF-MDCK (55.85 pN) (Fig. 3E black filled square), SF-MDCK (39.68 pN) (red filled upper triangle) and DF-MDCK (46.70 pN) (blue filled lower triangle) showed no significant differences between each other; however MDCK-MDCK interactions (sandal filled circle) showed substantially larger rupture forces (75.51 pN).

**Figure 3.**
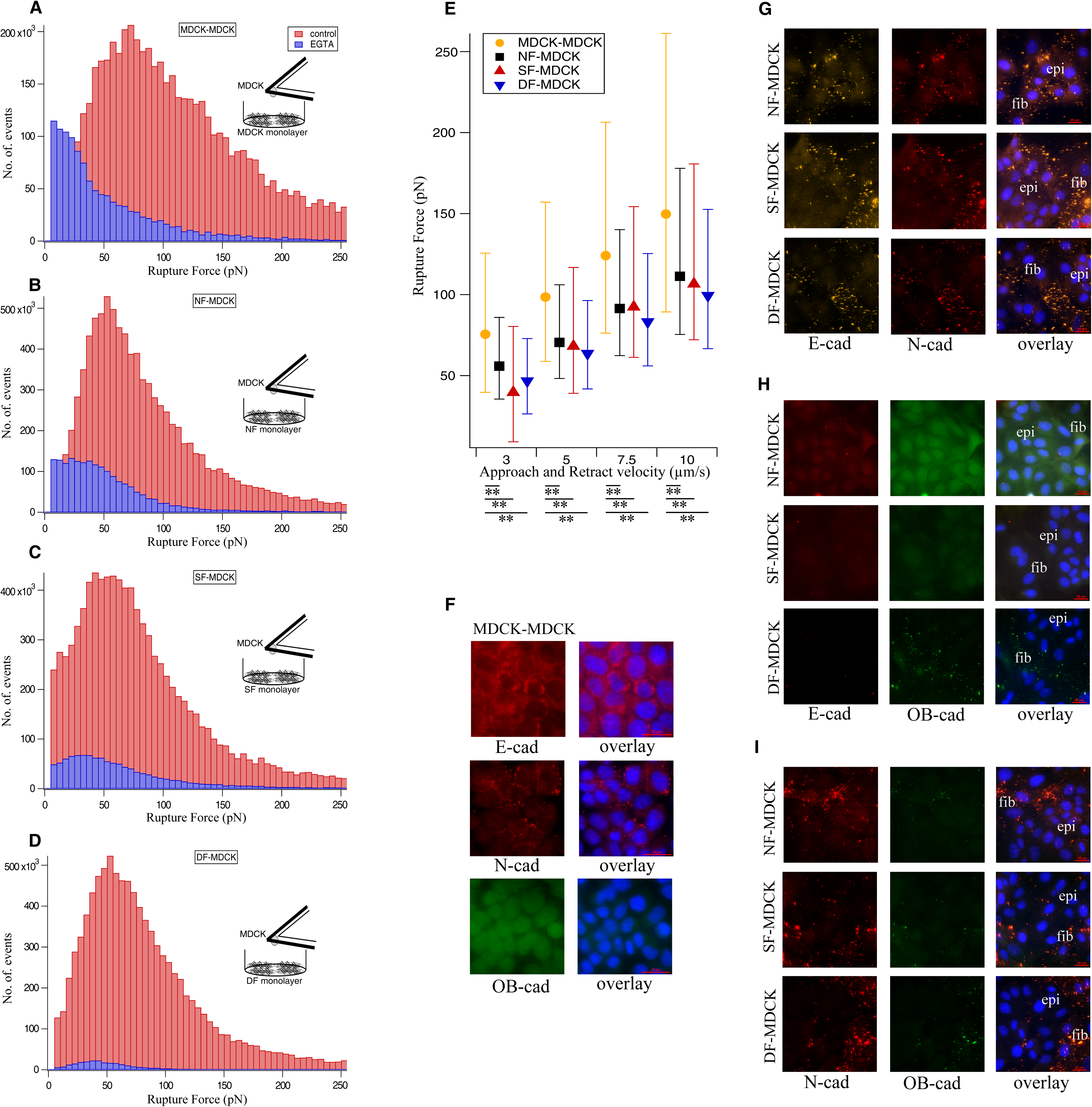
Heterocellular fibroblasts-MDCK interactions rupture force measurement and cadherin expression. Ruptures forces displayed as histograms recorded with (blue bar) and without EGTA (red bar) for (A) MDCK-MDCK, (B) NF-MDCK, (C) SF-MDCK and (D) DF-MDCK. (E) The cadherin rupture force shows linear relationship with cantilever approach and retract velocity for MDCK-MDCK (sandal circle), NF-MDCK (black filled square), SF-MDCK (red filled upper triangle), DF-MDCK (blue filled lower triangle). (F) Immunofluorescence data shows predominant E-cadherin expression in MDCK-MDCK adherens junction. Subpopulations of MDCK express N-cadherin but not OB-cadherin. Dual immunofluorescence data shows N-cadherin (red fluorescence in G, I) homophilic binding and loss of E-cadherin (H) in NF-MDCK, SF-MDCK and DF-MDCK. Due to the similar excitation and emission wavelength of fluorescence tags (secondary antibody), dual immunostaining (G) for N-cadherin (red fluorescence) and E-cadherin (sandal fluorescence) is difficult to interpret. The E-cadherin loss seen in (H) confirms the N-cadherin homophilic binding in (G). Only faint OB-cadherin expression (green fluorescence) observed in DF-MDCK (I). Blue fluorescence indicates DAPI-stained nuclei. Scale bar 20 μm.

We examined the distribution of different cadherin subtypes in MDCK-MDCK homo-cellular and fibroblast-MDCK hetero-cellular adherens junctions. MDCK monolayers were immuno-stained for E-, N- and OB-cadherin (Fig. 3F). We observed E-cadherin in the MDCK cell-cell junctions with a sub-population of MDCK cells expressing very little N-cadherin. In addition, there was no OB-cadherin expression in MDCK cells. To verify the cadherin expression in fibroblast-epithelial cell interaction sites, NF-MDCK, SF-MDCK and DF-MDCK were dual immuno-stained against the different cadherin subtypes (E/N, E/OB and N/OB). The secondary antibody with fluorescent tags that was used for detection of the primary anti-N-cadherin and anti-E-cadherin share almost the same excitation and emission wavelength. This made it difficult to differentiate between the E- and N-cad heterophilic interaction in NF-, SF- and DF-MDCK adherens junctions (Fig. 3G). Dual immunostaining for E- and OB-cadherins showed a reduction of E-cadherin and absence of OB-cadherin expression in co-cultures with NF-MDCK or SF-MDCK. The loss of E-cadherin was accompanied with faint expression of OB-cadherin in DF-MDCK cultures as well (Fig. 3H). In dual immunostaining for N- and OB-cadherins (Fig. 3I), only N-cadherin expression and no OB-cadherin expression were seen at the NF-MDCK and SF-MDCK and very little OB-cadherin at the DF-MDCK adherens junctions. The observation from these cadherin (E/OB and N/OB) subtypes helped to solve the E/N subtype issue and confirms the presence of N-cadherin in Fig. 3G. Control experiments showed no E-cadherin expression in NF-NF, SF-SF and DF-DF (Supplementary Fig. 3). In summary, the immunofluorescence data showed that N-cadherin is the predominant cadherin in the fibroblasts-MDCK adherens junctions. N-cadherin was exclusively seen in the fibroblasts-MDCK and not between MDCK-MDCK junctions in co-cultures. Initially, MDCK-MDCK interaction in MDCK cell cultures showed more E-cadherin and very little N-cadherin expression.

### Role of actin assembly in homo- and hetero-cellular adherens junctions

Cytochalasin D disrupts the actin assembly and results in cell softening (21). Here, we used 5 μM cytochalasin D to disassemble actin filaments to investigate the role of actin in both homo-cellular and hetero-cellular adherens junctions. As the drug was dissolved in DMSO, any effect of DMSO in cell-cell interaction had to be ruled out in control experiments before. The cityscape plot showed the rupture forces of cadherins bond rupture before and after the addition of cytochalasin D recorded from homo-cellular (Fig. 4A-D) and hetero-cellular (Fig. 4E-G) systems. The retract curves from control experiments (defined as no drug and no DMSO) (Supplementary Fig. 4A) or with DMSO (Supplementary Fig. 4B) showed no differences in the rupture patterns whereas with cytochalasin D (Supplementary Fig. 4C) dissimilar rupture events were observed. Treatment with cytochalasin D resulted in a reduction of the peak rupture force (Fig. 4 blue line) compared to DMSO addition (Fig. 4 red line) or control (Fig. 4 black line). The respective median rupture force values calculated for each condition for both homo-cellular and hetero-cellular junctions are listed in Table 2. The plot of median rupture forces (Supplementary Fig. 5) showed that the cadherin bond rupture force values were decreased in the presence of cytochalasin D for both homo- and hetero-cellular interactions; whereas, no significant differences were observed between control and DMSO rupture force values. This illustrates that the regulation of cadherin extracellular binding dynamics by intracellular actin filaments through their interaction with cadherin cytoplasmic domain.

**Table 2:**
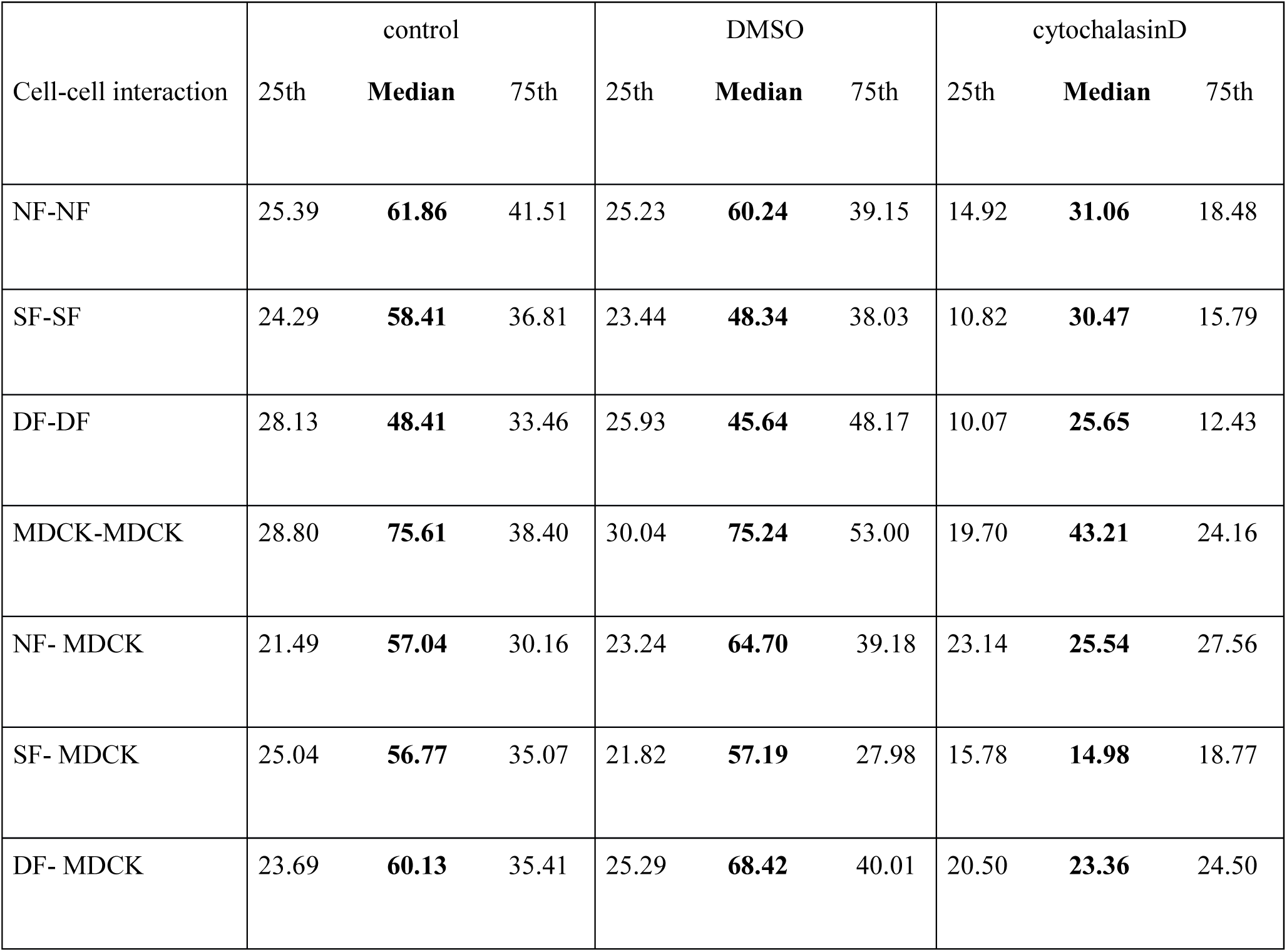
The median (bold values) rupture force (pN) values of cadherin mediated homocellular and heterocellular adherens junctions without drug (control), DMSO and cytochalasin D (5 μM).

**Figure 4.**
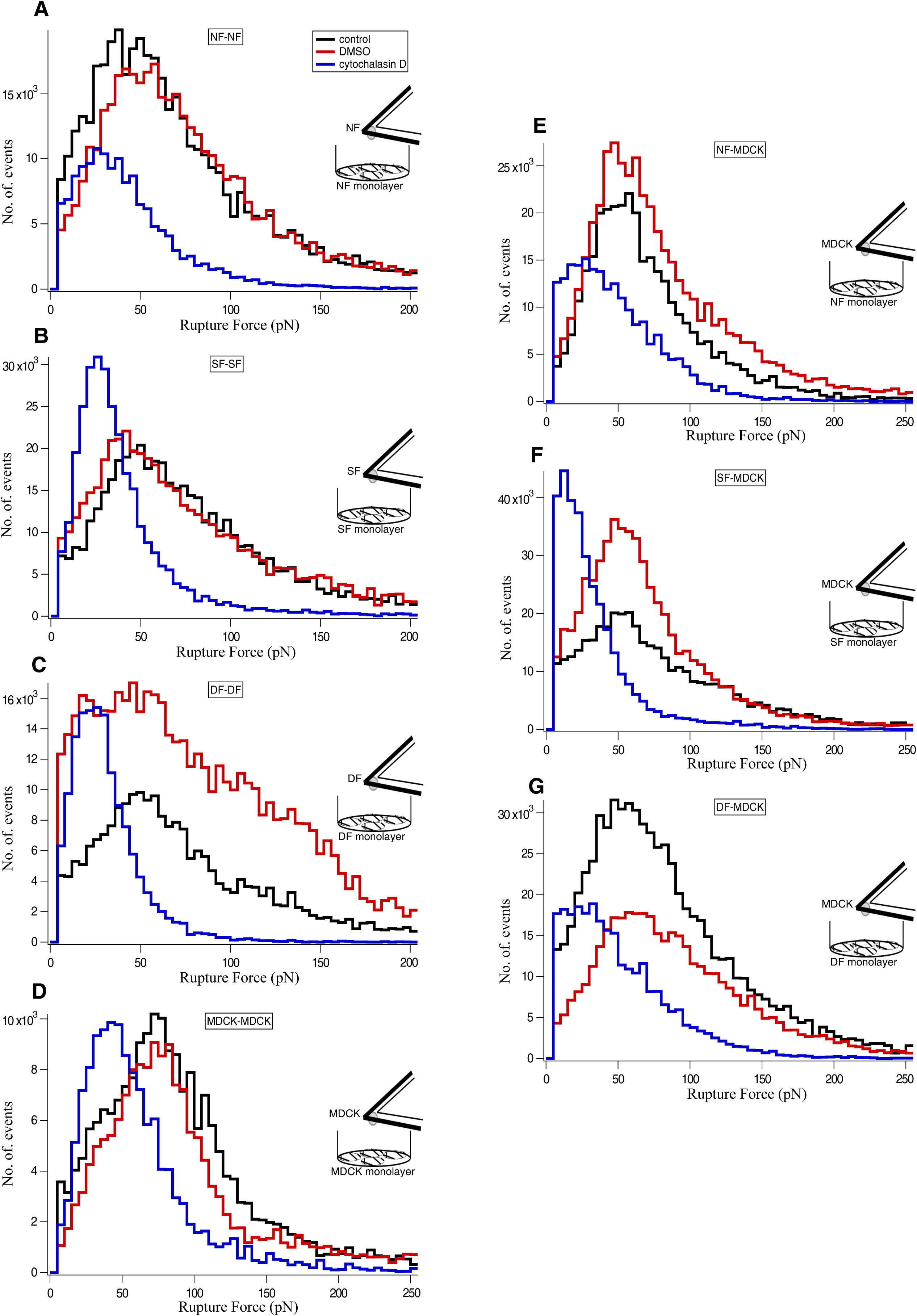
Effect of Cytochalasin D treatment on the rupture force of homocellular and heterocellular adherens junctions. The cityscape plot summarize the effect of cytochalasin D (5 μM) on homocellular- (A) NF-NF, (B) SF-SF, (C) DF-DF, (D) MDCK-MDCK and heterocellular- (E) NF-MDCK, (F) SF-MDCK, (G) DF-MDCK adherens junctions. Rupture events were recorded without the drug as control (black line), with DMSO (red line) and with cytochalasin D (blue line). The corresponding median values were listed in Table 2 and plotted in Supplementary Fig. 5.

### Myosin inactivation strengthens the N-/OB-cadherin heterophilic binding

ML-7 inhibits the MLCK activity by inhibition of myosin cross-linking to the actin filaments. As a consequence, there is reduction in contractile stresses, which leads to a softening of the intracellular actin cytoskeleton (23). Here we used 5 μM ML-7 to inhibit the myosin activity and determined the role of myosin in both homo-cellular and hetero-cellular adherens junctions. The cityscape plot showed the rupture forces of cadherin bond ruptures before and after the addition of ML-7 recorded from homo-cellular (Fig. 5A-D) and hetero-cellular (Fig. 5E-G) systems. The retract curves from control (no drug and no DMSO) (Supplementary Fig. 6A), DMSO (Supplementary Fig. 6B) and with ML-7 (Supplementary Fig. 6C) showed no difference in the rupture patterns except for DF-DF. No significant shift in rupture force peaks was observed in control (Fig. 5 black line), DMSO (Fig. 5 red line) and ML-7 (Fig. 5 blue line). Only DF-DF showed an increase in the rupture force after treatment with ML-7 (Fig. 5C blue line). Comparing the retract curves of controls (Supplementary Fig. 7A), DMSO (Supplementary Fig. 7B) or ML-7 (Supplementary Fig. 7C) treatment of different fibroblast cultures, DF-DF showed distinctive large rupture events in ML-7 treated force curves. The respective median rupture force values calculated for each condition for both homo-cellular and hetero-cellular junctions were listed in Table 3. The median plot (Supplementary Fig. 8A, B, D, E, F, and G) showed no significant change in the rupture force values for cell-cell interactions in NF-NF or SF-SF cultures in the presence of ML-7 comparing to that of control and DMSO. In case of DF-DF, the median rupture force value (Supplementary Fig. 8C) increased significantly after the addition of ML-7. Interestingly, no significant differences were observed in control and DMSO rupture force values in DF-DF. Possibly, the intracellular myosin inactivation by MLCK inhibition does not affect the cadherin homophilic binding. In case of DF-DF which express N-cadherin/OB-cadherin heterophilic binding, myosin inactivity seems to strengthen heterophilic cadherin interactions.

**Table 3:** The median (bold values) rupture force (pN) values of cadherin mediated homocellular and heterocellular adherens junctions without drug (control), DMSO and ML-7 (5 μM).

| Cell-cell interaction | control |  |  | DMSO |  |  | ML-7 |  |  |
| --- | --- | --- | --- | --- | --- | --- | --- | --- | --- |
|  | 25th | **Median** | 75th | 25th | **Median** | 75th | 25th | **Median** | 75th |
| NF-NF | 24.48 | **58.81** | 45.80 | 19.67 | **52.04** | 38.84 | 16.49 | **48.87** | 33.25 |
| SF-SF | 27.37 | **66.20** | 38.17 | 34.66 | **53.12** | 41.96 | 28.72 | **48.98** | 36.52 |
| DF-DF | 24.29 | **53.19** | 34.65 | 23.19 | **45.49** | 31.87 | 48.01 | **99.69** | 75.06 |
| MDCK-MDCK | 24.53 | **69.71** | 43.41 | 27.11 | **67.22** | 56.11 | 26.03 | **69.69** | 42.50 |
| NF- MDCK | 18.58 | **52.06** | 25.22 | 17.71 | **54.74** | 23.17 | 17.82 | **54.41** | 22.91 |
| SF- MDCK | 17.42 | **48.95** | 16.58 | 15.51 | **53.87** | 17.37 | 15.05 | **41.44** | 16.73 |
| DF- MDCK | 11.99 | **48.35** | 18.55 | 12.98 | **47.05** | 18.15 | 15.24 | **52.32** | 23.10 |

**Figure 5.**
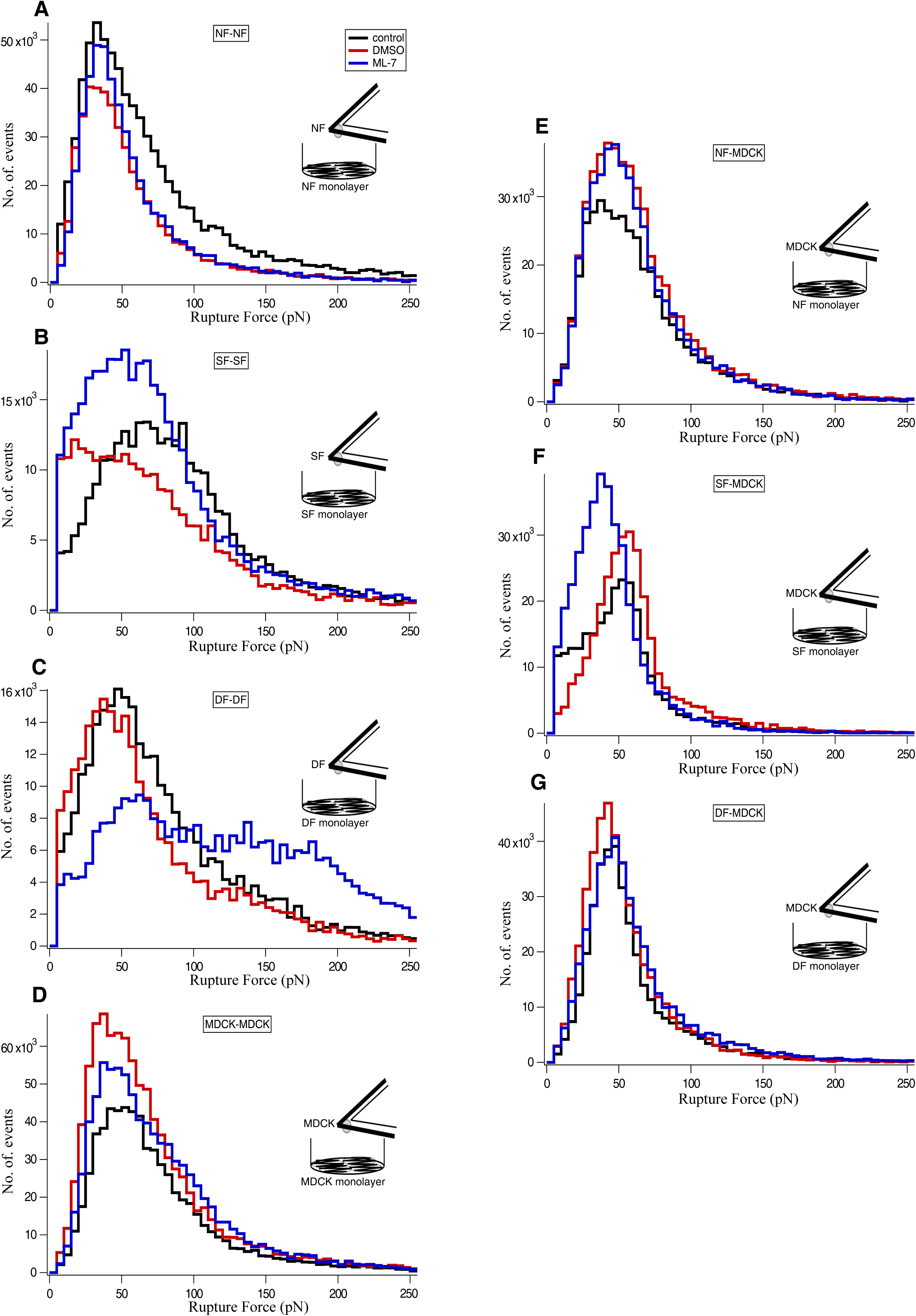
Effects of ML-7 treatment on the rupture force of homocellular and heterocellular adherens junctions. The cityscape plot summarize the effect of ML-7 (5 μM) on homocellular- (A) NF-NF, (B) SF-SF, (C) DF-DF, (D) MDCK-MDCK and heterocellular- (E) NF-MDCK, (F) SF-MDCK, (G) DF-MDCK adherens junctions. Rupture events were recorded without the drug as control (black line), with DMSO (red line) and with ML-7 (blue line). The corresponding median values were listed in Table 3 and plotted in Supplementary Fig. 8.

## Discussion

In this work, we performed AFM-based SCFS on homo- and hetero-cellular arrangements between different fibroblasts (NF, SF and DF) and MDCK cells. We measured the homophilic and heterophilic cadherin adhesion rupture forces using AFM-SCFS. Immunofluorescence staining allowed us to visualize the presence of such homophilic and heterophilic pairs of E, N and OB cadherins. Our results showed that homophilic adhesions were stronger than heterophilic adhesions. In addition, our results suggest a role of the intracellular actin cytoskeleton in homophilic and heterophilic cadherin bonds modulating extracellular binding strength. With differing binding capacity and specificity, cadherins of cellular adherens junction play an important role in intra- and inter-cellular mechano-signalling for force transmission. So far, AFM and optical tweezers based SMFS explored the binding strength and kinetics of various cadherin types - both homophilic and heterophilic binding (14, 24, 25, 26). Most of these studies were carried out with cadherins which were overexpressed or purified and lacking certain domains (for example, recombinant constructs lacking the cytoplasmic domain). Here, we used AFM based SCFS to measure the rupture forces of cadherin-cadherin bonds in or between cells, i.e. the cadherins measured were in their native state. This setup allowed us to study the cadherin pair formation and their bond rupture forces in physiologically relevant homo- and hetero-cellular arrangements. Previously, using this setup VE-, E- and N-cadherin homophilic pair formation, their binding strength and kinetics were studied using the homo-cellular arrangements made with HUVEC, L-M (TK-) and CHO cells, respectively (27, 28). In a similar way, we attached different types of fibroblasts (NF, SF and DF) to the AFM cantilevers and put them into contact with monolayers of the same type of fibroblasts. The measured rupture forces showed that NF-NF and SF-SF exhibit stronger interactions than DF-DF. Immunofluorescence studies revealed that N/N-cadherin homophilic pairs enforced the inter-cellular adherens junctions in NF and SF. Whereas in the case of DF, N-/OB-cadherin heterophilic pairs were seen in the cellular adherens junctions and this contributes to their weaker interaction. NF and SF were shown to express α-sma, but no large stress fibres. Thus both cell types were considered as fibroblast phenotype (29). When SF was seeded in a physiological environment such as a decellularized dermal matrix, cells expressed large stress fibres and thus showed a proto-myofibroblast or myofibroblast phenotype (30). In contrast, DF showed α-sma positive large stress fibres and thus were considered as myofibroblast phenotype (29). In comparison to earlier reports (1, 14), N-cadherin expression was seen in the normal fibroblast phenotype (in this study: NF and SF) and OB-cadherin expression in the myofibroblast phenotype (DF). In contrast to rat fibroblasts, which show a transition in expression from N-to OB-cadherin triggered by TGF-β1 (14), we found in our study that DF expressed both N- and OB-cadherins and N/OB heterophilic binding. Fibroblasts extracted from the palmar region (cords and nodules) of patients with Dupuytren’s disease express stress fibres and thus exhibiting a myofibroblastic phenotype (29) without any mechanical or biochemical stimulation such as TGF-β1 (14). Biochemical expression of N-cadherin was observed in Dupuytren’s myofibroblast and results from a collagen gel contraction study showed that myofibroblasts displayed reduced contraction in the presence of N-cadherin blocking peptide (1). Obviously, N-cadherin has an important function in myofibroblast intercellular adherens junctions. By immunofluorescence, N-cadherin and OB-cadherin expression and their homophilic (in NF and SF) and heterophilic (in DF) pair formation were observed in all three fibroblast phenotypes. In this study, we found that the presence of different cadherins was strongly correlated with the rupture forces measured by SCFS.

In our study, the E/E-cadherin homophilic interactions in MDCK showed rupture force values closely related to previous studies (28). This confirms the initial adhesion in MDCK homo-cellular arrangements could be largely dominated by E-cadherin homophilic binding that displays the larger rupture force. As previously shown, in contrast to primary epithelial cell that do not express the N-cadherin subtype, MDCK sub-populations such as trypsin sensitive MDCK are characterized by N-cadherin expression (16) which we could confirm the N-cadherin expression seen in MDCK cultures with our immunofluorescence analyses.

In tissue and sub-tissue level biology, multicellular interactions are orchestrated through various cell-cell junction mechanisms which, in turn, coordinate individual cell type actions such as directed cellular migration and wound contraction. As central component of the adherens junction, the cadherin transmembrane domain plays a key role in force transmission between the intracellular environment of different cell types and the intercellular space through cadherin type binding specificity. This phenomenon could have implications for biological analytical methods, e.g. for cell sorting. In response to tissue injury and during wound healing, direct contact between epithelial cells and underlying fibroblasts modulate the expression levels of key enzymes such as matrix metalloproteinase-2 and -9, which are important for the wound healing process (31). Through activation by the cytokine TGF-β1 in the ECM or by mechanical injury of epithelial cells, the biochemical expression of α-sma and type I and III collagen was induced in co-cultured fibroblasts (32). These observations brought the knowledge of investigating the adhesion proteins involved in hetero-cellular interactions such as epithelial cell-fibroblasts interaction. For SCFS hetero-cellular studies, MDCK cells were attached on the cantilever and brought in contact with fibroblasts grown in monolayers in a Petri dish. This experimental set-up was chosen to measure the rupture forces of the N/N homophilic and E/N, E/OB and N/OB heterophilic bonds. Distinct peaks were not observed in the rupture force histograms, but the observed heterophilic bond results are able to be discussed with previous results. For example, an earlier SCFS study did not show any occurrence of heterophilic interactions between E-cadherin and N-cadherin (28). Contrarily, a single molecule study shows the existence of such E/N cadherin heterophilic interactions. Presumably, shorter contact/dwell times (millisecond) used in the former studies (33) could be the reason for not recognizing heterophilic interactions as found in our study. In standard experimental settings, shorter contact times between the cells in the petri dish and on the cantilever were used to prevent nonspecific binding. Deliberately, we chose a different experimental design with longer contact time of 2 sec in the SCFS setup which enabled us to follow both homophilic and heterophilic cadherin interactions. Despite of the changed protocol, distinct peaks could not be resolved in the histograms. This might be due to the possibility that N/N and E/N rupture forces share similar values. In case of DF-MDCK, no distinct peaks of E/OB and N/OB were seen which could be due to similar rupture forces. This leads to the question if single molecule kinetic studies using AFM or optical tweezers are suitable to measure homophilic and heterophilic cadherin pairs with definite set of contact times.

E-/N-cadherin heterotypic adhesion sites reinforced by local cytoskeletal reorganization were observed between IAR-2 epithelial cells and RAT-1 fibroblasts using immunofluorescence staining (34). This mechanically active heterotypic contact between N-cadherin expressing cancer associated fibroblasts and an E-cadherin expressing epithelial (A431) cancer cell line (A431) enables fibroblasts to steer cancer cell invasion (18). Loss of E-cadherin was observed in co-cultures of fibroblast with epithelial cells, whereas normal fibroblasts can induce E-cadherin loss to promote EMT in gastric cancer (20). In chronic inflammatory conditions, epithelial cell-fibroblast interactions stimulate EMT in human bronchial epithelial cells from chronic obstructive pulmonary patients (19). Accordingly, we found reduced E-cadherin and increased N-cadherin in our multi-cell cultures with immunofluorescence which might imply the initiation of an EMT process. Furthermore, N/N homophilic adhesion (NF-MDCK and SF-MDCK) and N/OB heterophilic adhesion (DF-MDCK) were present at the interaction sites between epithelial cells and fibroblasts.

In AFM based SCFS, a varying cantilever pulling rate allowed for characterizing the cadherin binding strength. Rupture forces generally increase with increasing pulling rate, which leads to increased loading rates (35). In this study, E-cadherin and N-cadherin homophilic and OB-cadherin heterophilic binding rupture forces showed a linear relationship related to the pulling rate. In the fibroblast homo-cellular arrangement, N-cadherin homophilic binding was stronger in NF and SF compared to N/OB-cadherin heterophilic binding in DF. Similarly, in fibroblast-epithelial cell hetero-cellular arrangement, all three fibroblast types interacting with MDCK show similar rupture forces. In general, E-cadherin homophilic binding in MDCK homo-cellular arrangement displayed the strongest binding strength which reflects previous findings (14, 27, 28).

Differences in force peak values can be found when results are compared to other studies. Due to the stochastic process of cadherin, protein binding forces can be distributed differentially. Rupturing of molecular bonds is always effected by thermal fluctuations, leading to varying rupture forces and thus cadherin binding events are stochastic (36). Even the VE-, N- and OB-cadherin SMFS and SCFS measurements showed three different interaction forces, as the three force peaks were present in rupture force histograms (14, 24). However, cadherin pairs (VE-, E- and N-) exhibited single force states as well which correlates well to results found in earlier SCFS studies (27). Similarly, we observed one single force peak in the histograms which correspond to a single rupture force of cadherin bond unbinding.

Cell-cell adhesion is mediated by cadherins in adherens junctions. Cadherins are linked with their cytoplasmic domain to the intracellular actin cytoskeleton through adaptor proteins such as α- and β-catenin (37). Disruption of actin filaments by cytochalasin D affected the cadherin extracellular domain homophilic and heterophilic binding dynamics in our study. It seems that the inactivation of actin filaments with cytochalasin D has a direct effect on the cadherin extracellular binding activity by altering the cadherin cytoplasmic link to the actin filaments (14). However, this phenomenon was found exclusively for OB-cadherin homophilic binding (14). In our study we could show a similar effect for both homophilic (N/N and E/E) and heterophilic (E/N, N/OB and E/OB) adhesion in homo- and hetero-cellular arrangements.

ML-7 inhibits the activity of MLCK by interacting with the phosphorylation event of myosin light chain (MLC). Thus the binding of myosin to actin filaments and ATPase driven contractile force generation are inhibited (38, 39, 40). In the current study, disabling actin-myosin contraction using ML-7 showed no effect on the cadherin extracellular binding dynamics except for N-/OB-cadherin heterophilic binding. Myosin inactivation particularly strengthened the N-/OB-cadherin extracellular binding activity demonstrated by the change of rupture forces. A hypothetical biophysical mechanistic pathway that could explain the observed N/OB-cadherin reinforcement effect is stated in Fig. 6. Myosin II acts as an actin crosslinker (41) whereas myosin VI acts as a mediator protein, which binds cadherin to actin filaments (42). Loss of myosin II selectively inhibits myofibroblast differentiation in fibroblasts of fibrotic lung when compared to healthy phenotype (43). From our current findings and previous results from others, we speculate that (Fig. 6): (1) There is no influence of actomyosin contraction or inactivated myosin on homophilic or heterophilic cadherin extracellular binding dynamics (excluding N/OB); (2) myosin is creating tension in actin filament network, which weakens the N/OB-cadherin heterophilic bond, while inactivation of myosin strengthens this bond; (3) myosin inactivation enhances the N/OB-cadherin reinforcement by the detachment from the cadherin-actin complex. (4) As a consequence, actin filaments per se reinforce and stabilize the cadherin extracellular binding. Draw-backs of the current study include the analysis of biochemical expression levels of all myosin types (1-6) and respective localization associated with other functional abilities such as anchoring cadherin-catenin complex to the actin filaments in the cell-cell adhesion sites. Investigations into downstream intracellular signalling pathways are necessary to study further details on the involvement of other signalling molecules (adaptor proteins) in cadherin homophilic and heterophilic adhesion.

**Figure 6.**
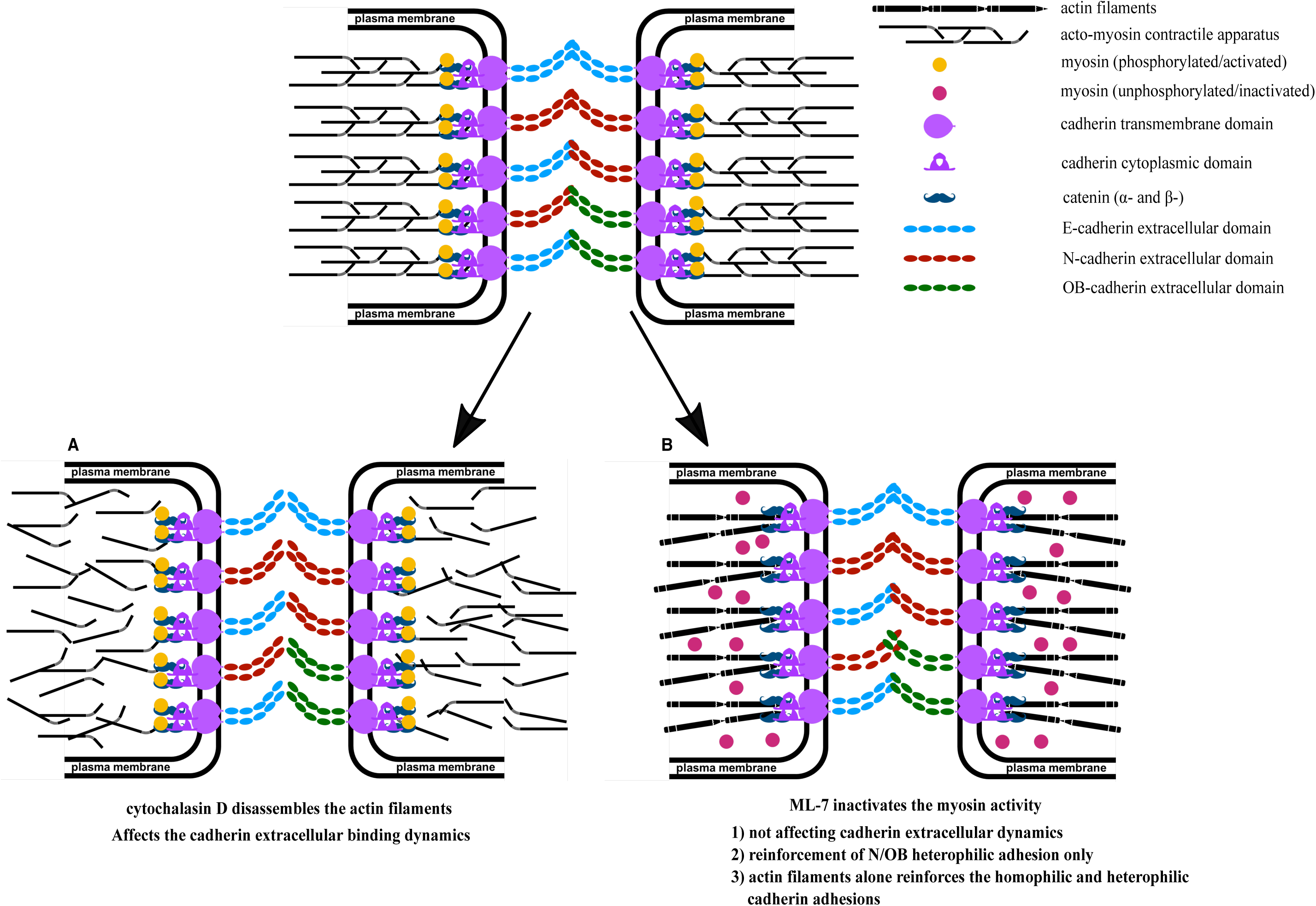
Acto-myosin contractility influences cadherin extracellular domain binding dynamics. Cartoon representations depict the effect of actin filaments disruption (A) and myosin inactivation (B) on cadherin homophilic and heterophilic adhesion pairs. (A) Actin filaments disassembly resulting of cytochalasin D treatment leads to weakening of homophilic and heterophilic cadherin adhesions. (B) Myosin inactivation by inhibiting myosin light chain kinase (MLCK) using ML-7 treatment leads to N/OB heterophilic adhesion reinforcement whereas dissimilar effects were seen in other cadherin homophilic and heterophilic adhesions. This pictures the stabilization and reinforcement of cadherin homophilic and heterophilic adhesion by actin filaments, with no cross linkers-myosin generating contractile forces and with myosin inactivation at the cadherin-catenin-actin complex.

## Conclusions

So, in understanding the biophysical dynamics of cell-cell adhesion, the underlying actin/adherens junctions and its associated proteins have also to be considered. Our findings showed that myosin inactivation provides mechanical strengthening of N/OB heterophilic adhesion and dissimilar effects on other homophilic and heterophilic adhesion. Based on our results, further studies are required to investigate the multifunctional role of myosin types, actin filaments and other associated proteins in cell-cell adhesion. SCFS can be a suitable experimental setting to examine the role of intracellular proteins involved in various cellular processes, specifically cell-ECM adhesion and here cell-cell adhesion, if one can design the experiments accordingly.

## Materials and Methods

### Cell culture

Cell culture was performed as described previously (29). Fibroblasts were harvested from tissues of patients undergoing hand surgery (approved by the local Ethics Committee-Ärztekammer Bremen, #336/2012) and isolated as described previously (29). Cells were grown until the passage-9 for fibroblasts and 13 for MDCK in DMEM medium and incubated at 37°C in a humidified atmosphere of 95% air and 5% CO_2_. Cell culture was established for two days before proceeding with further SCFS measurements. Medium was supplemented with 10% fetal bovine serum (FBS) and 2% penicillin-streptomycin.

### Cantilever functionalization

The silicon-nitride tipless cantilevers (Nanoprobe SPM Tips, NP-OW 9861) were washed with 1 % SDS (sodium dodecyl sulphate), Helizyme (B. Braun Vet Care GmbH) and distilled water solution each for overnight. The cantilevers were then treated with plasma (Ar) at high power for 5 min. In order to functionalize the plasma treated cantilevers with concanavalin A (conA) (C2010, Sigma-Aldrich), the cantilevers were placed in a phosphate buffered saline (PBS) solution containing conA (2 mg/ml) for 2 h at room temperature. The conA coated cantilevers were stored in PBS at 4°C (44).

### Cell attachment to the cantilever

Prior to cell-cell adhesion measurements, cells that were used for attachment to the cantilever were released from the culture flask by treatment with trypsin for 2 min and trypsin was neutralized by centrifugation and replenishment with new medium. The trypsinized cells were transferred into the Petri dish containing firmly attached cell monolayers that are grown for two days. After 5 min incubation at 37°C, the Petri dish was used for the single-cell force spectroscopy-AFM setup. The conA functionalized cantilever was then placed over a suitable cell with round morphology which initiated its attachment to the cell monolayer. Then, the conA coated cantilever was approached towards the cell with a 3.5 nN maximum loading force for 5 sec at a velocity 5 μm/sec until the cell was captured. The cantilever with attached cell was taken few μm away from the cell monolayer and the whole setup was left undisturbed for 10 min in order to establish firm cell adhesion to the cantilever.

### AFM cell adhesion force measurements and data analysis

Single-cell experiments were conducted using a MFP3D AFM (Asylum Research, Santa Barbara, CA, USA). An optical microscope (Zeiss Axiovert 135, Zeiss, Oberkochen) was combined with the AFM to be able to control cantilever and sample positioning. All measurements were performed with 15 tipless cantilevers with a nominal spring constant 60 pN/nm. The Petri dishes with the cell monolayer were fixed to an aluminium holder with vacuum grease and mounted on the AFM stage with two magnets. The AFM head including the sample was enclosed in a homebuilt polymethacrylate (PMMA) box in order to inject and maintain 5% CO_2_. Force maps were recorded on cell monolayer to obtain cell-cell rupture force. First, the spring constant of the conA coated cantilever was calibrated by using the thermal tune method on a cleaned and stiff surface (45) and then cell capturing followed by cell-cell adhesion force curves were recorded. For force curves, we used typically a maximum loading force of 3nN with 2 s dwell time at a velocity (approach and retract) of 3 μm/sec.

The data analysis package IGOR (wave metrics, Lake Oswego, OR, USA) was used to evaluate the rupture force from the retract force curve. The retract curve contains two different patterns - jumps and tethers. Jumps in the retract curve correspond to the rupture of cadherin bonds, whereas plateaus correspond to pulling of membrane tethers. The height of all jumps was multiplied to the cantilever spring constant in order to obtain the rupture force. By changing the approach and retract velocity (5, 7.5 and 10 μm/sec), we measured the loading rate dependent rupture forces within the cadherin bonds. Rupture forces calculated from all rupture events were presented in histograms. Each category of experiments was repeated two to four times (n=2 to 4). For each category, 30 to 40 force maps (one force map contains 24 force curves) were analyzed.

### EGTA, Cytochalasin D and ML-7 addition

For demonstrating Ca^2+^ specific cell-cell interactions, control experiments were performed with 7.5 mM EGTA (Sigma-Aldrich). For drug induced changes on cell-cell adhesion measurements, cytochalasin D (C8273, Sigma-Aldrich) and ML-7 (I2764, Sigma-Aldrich) were used at 5 μM working concentration. Substances were solubilized in DMSO to a stock solution of 200 μM. From this stock solution, 100 μL were added to cultures to a final concentration of 5 μM. To exclude the nonspecific effects of DMSO, control experiments with DMSO were performed in parallel and plotted with drug induced changes in cell-cell adhesion.

### Immunofluorescence staining

Regarding immunofluorescence experiment for fibroblast-epithelial cell interaction, co-culturing of fibroblast and epithelial cells was performed in 1:2 ratio. Two days after seeding, cells were fixed with 3.7% formaldehyde for 15 min and permeabilized with 0.1% Triton X100 for 3 min. Samples were washed with PBS after each step and blocked with 3 % goat serum and then incubated with primary antibodies, anti-N-cadherin 1:200 dilution (rabbit polyclonal; sc-7939, Santa Cruz Biotechnology), anti-OB-cadherin 1:50 dilution (mouse monoclonal; sc-365867, Santa Cruz Biotechnology) and anti-E-cadherin 1:50 dilution (goat polyclonal; AF748, R&D systems) at 4°C overnight. After incubation, samples were washed with PBS containing goat serum. Then samples were blocked with 3 % goat serum and then incubated with respective secondary antibodies, e.g. cy3 anti-rabbit IgG (711-165-152, Jackson ImmunoResearch Laboratories, Inc.) at 1:200 dilution, FITC anti-mouse IgM (315-095-020, Jackson ImmunoResearch Laboratories, Inc.) at 1:100 dilution and Rhodamine/TRITC anti-goat IgG (305-025-045, Jackson ImmunoResearch Laboratories, Inc.) at 1:100 in a dark environment. For multicolor staining (dual staining), a sequential (staining one protein after another) incubation of primary and secondary antibodies was performed. Then samples were washed with PBS and stored with ProLong Diamond Antifade Mountant with DAPI (P36966, ThermoFisher Scientific) at 4°C prior to image acquisition. The cells were visualized with a 100x oil-immersion objective mounted on Nikon Eclipse Ti Inverted epifluorescence Microscope (Nikon Instruments Inc., Melville, New York).

### Statistical analysis

Statistical differences for the median values of rupture force of cadherins present in homo-cellular and hetero-cellular systems of the AFM measurements were determined by Wilcoxon test, calculated in IGOR software. * and ** indicate statistically significant differences for p-values < 0.05 and p < 0.005, respectively.

## Acknowledgements

We thank Holger Doschke for developing the data acquisition and analysis software and also for helpful discussions. We thank Prof. Dorothea Brüggemann for providing the fluorescence microscope. We also thank Dr. Mario Waespy and Christin Goldbaum for their fruitful discussions. This work was supported by the DFG grant MA4172/12-1.

## Data availability statement

All datasets generated during and/or analysed during the current study are available from the corresponding author.

## Author contributions

PK designed and performed the research and was also performing AFM experiments, fluorescence experiments, data analysis and manuscript preparation. UM provided the different fibroblast types, analytical material and substances and, was involved in the manuscript preparation. GB isolated and provided the cells and also provided antibodies. MR designed the experimental setting and was involved in data acquisition, data analysis and preparation of the manuscript.

