## Supplementary information for "Homophilic and heterophilic cadherin bond rupture forces in homo- or hetero-cellular systems measured by AFM based SCFS"

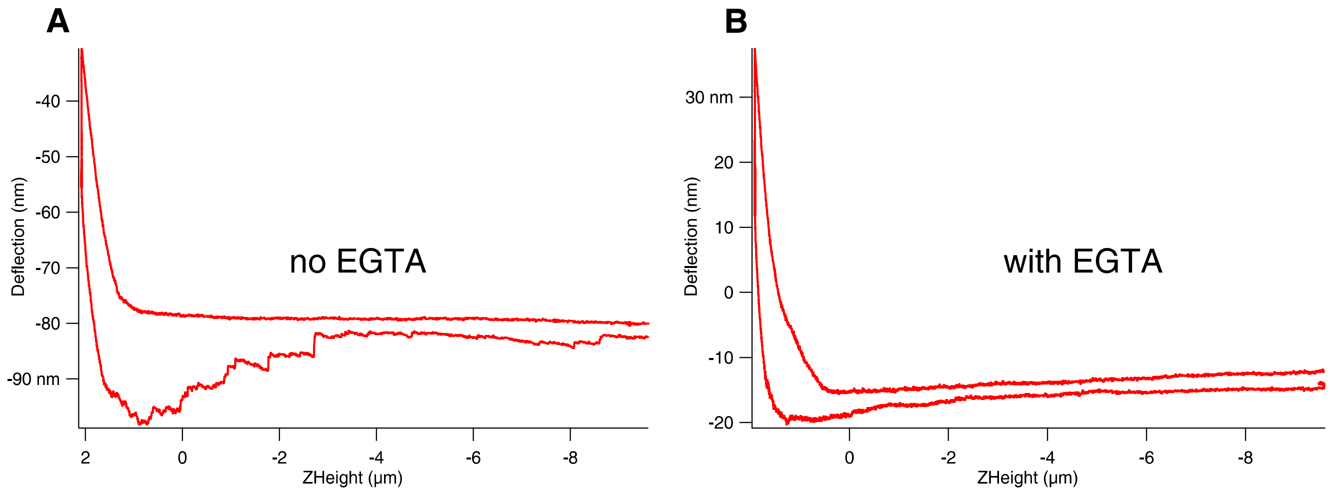


**Supplementary figure 1** Force curves obtained during cell-cell interaction (here MDCK-MDCK) show distinct rupture events under normal conditions (no EGTA) (A), which in the presence of EGTA, corresponding to low or no $Ca^{2+}$ present, disappear (B).

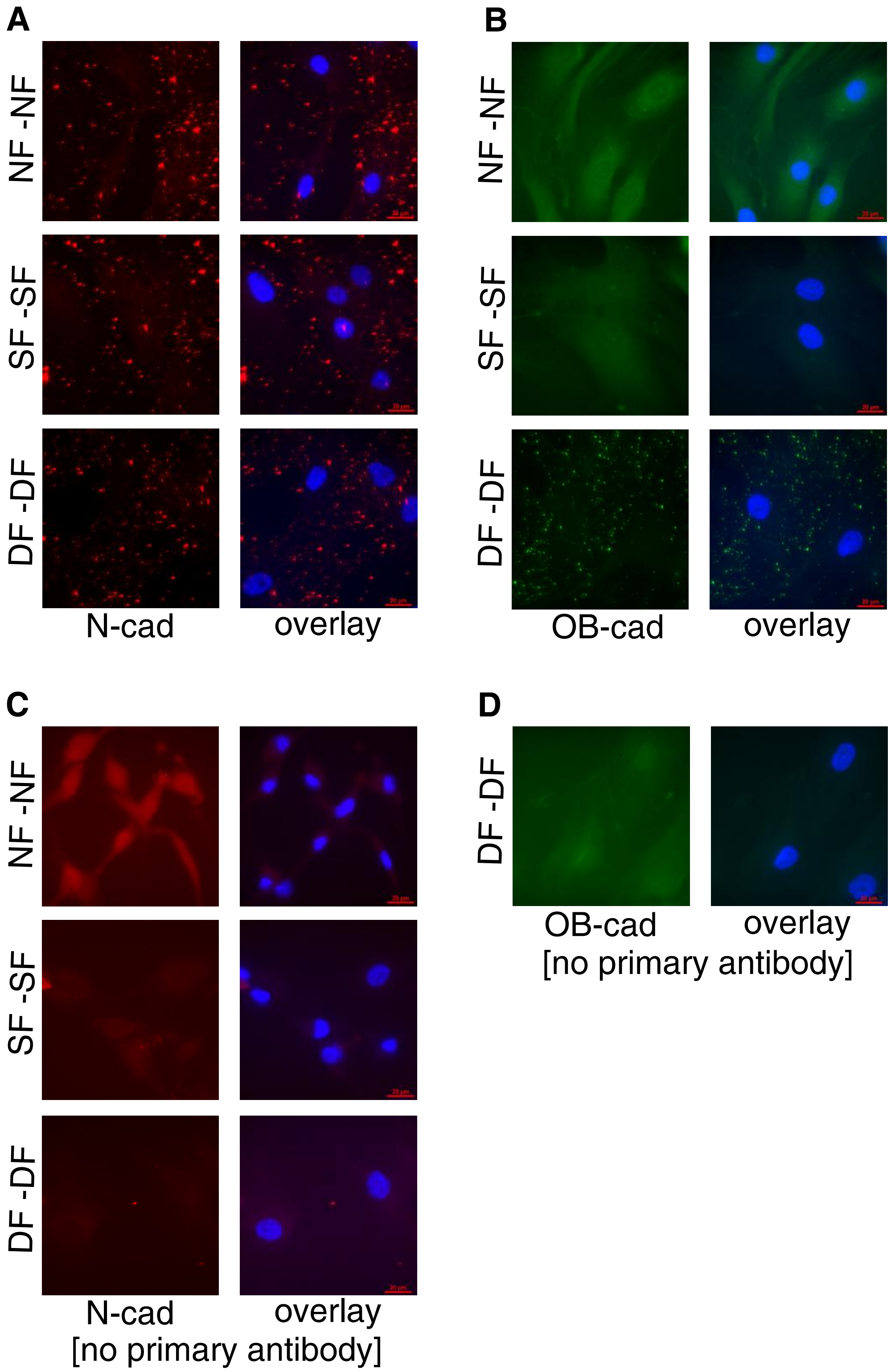


**Supplementary figure 2** Immunostaining of fibroblasts adherens junctions for N- and OB- cadherin shows N-cad expression (red fluorescence) in all fibroblasts (A) and OB-cad expression (green fluorescence) only in DF interaction sites (B). In the right column (overlay) DAPI staining of the nucleus is overlayed with the corresponding antibody staining. In the control measuremnts (C&D) unspecific binding of the secondary antibody was checked by staining without the corresponding primary antibody for N-cad (C) and OB-cad (D). Only a weak homogenous background fluorescence signal was detected showing that the secondary antibody specifically binds the primary antibody. Scale bar 20 µm.

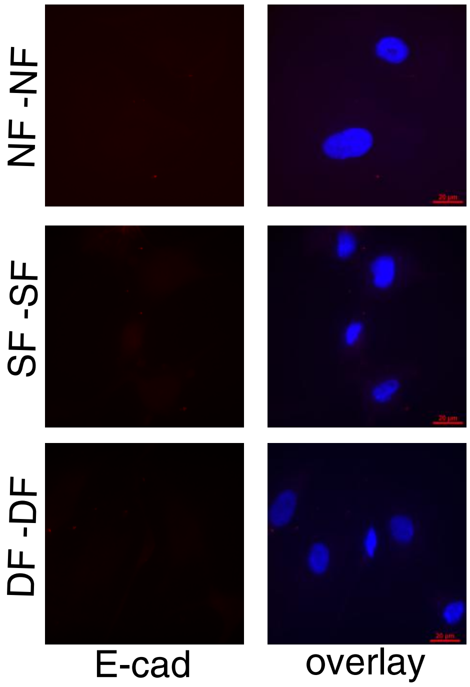


**Supplementary figure 3** Immunostaining of fibroblasts adherens junctions for E-cadherin shows no expression of E-cad in the NF-NF, SF-SF and DF-DF interaction sites. Scale bar 20 µm.

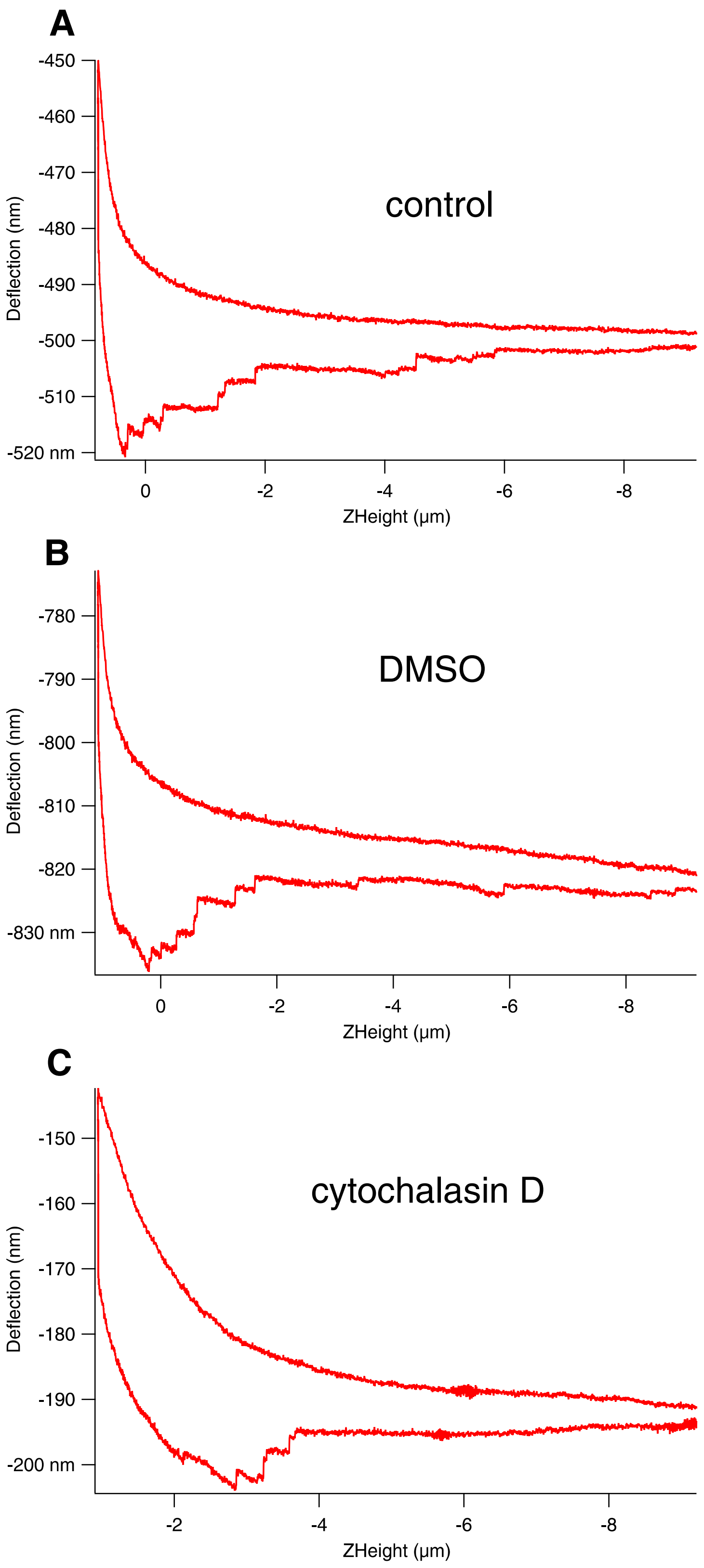


**Supplementary figure 4** Force curves measured during cell-cell interaction (here for NF-NF) shows rupture events in all three experimental conditions: control in normal DMEM medium (A), medium plus DMSO (B), and medium with cytochalasin D (C).

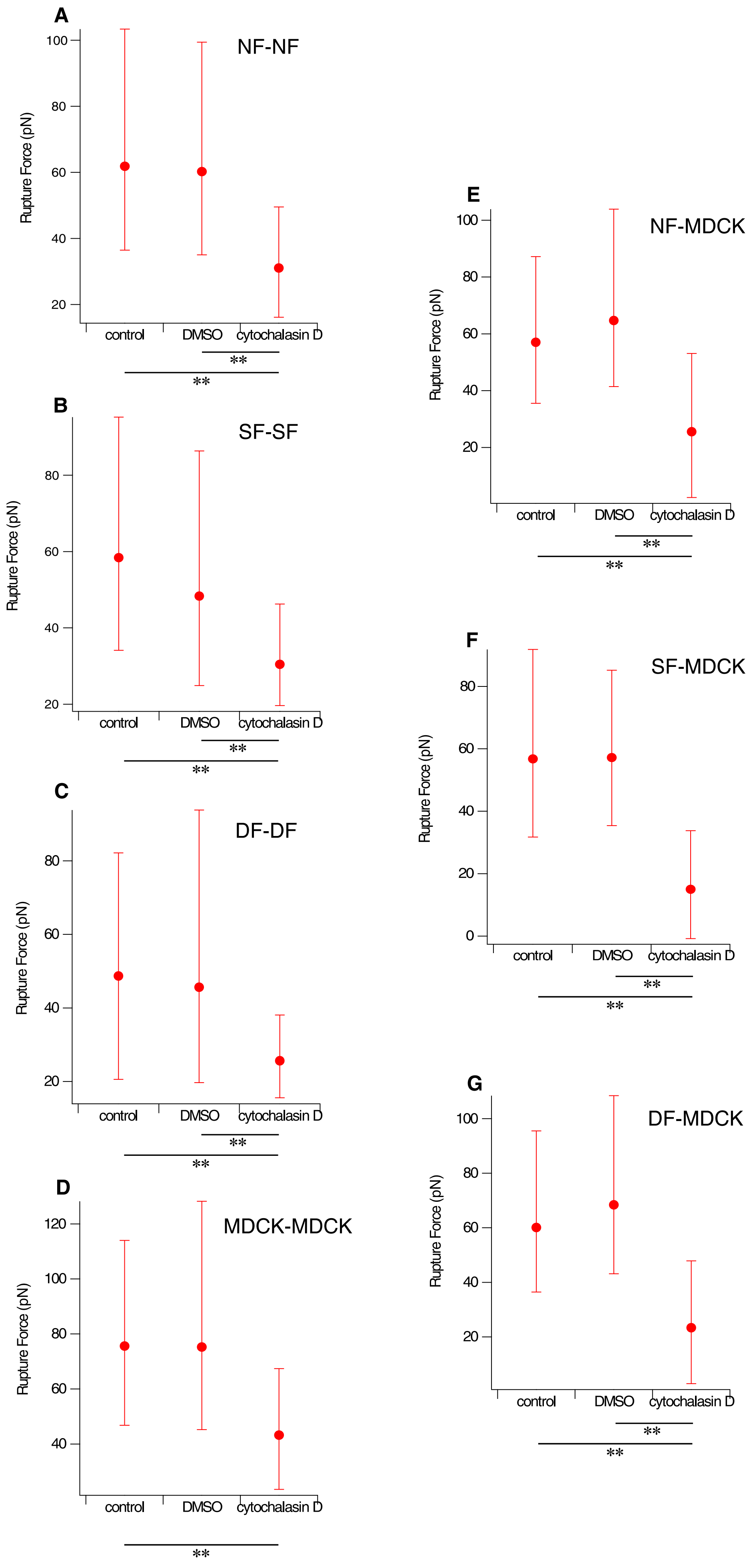

A
NF-NF
Rupture Force (pN)
100
80
60
40
20
control
DMSO
cytochalasin D
**
**
B
SF-SF
Rupture Force (pN)
80
60
40
20
control
DMSO
cytochalasin D
**
**
C
DF-DF
Rupture Force (pN)
80
60
40
20
control
DMSO
cytochalasin D
**
**
D
MDCK-MDCK
Rupture Force (pN)
120
100
80
60
40
control
DMSO
cytochalasin D
**
E
NF-MDCK
Rupture Force (pN)
100
80
60
40
20
control
DMSO
cytochalasin D
**
**
F
SF-MDCK
Rupture Force (pN)
80
60
40
20
0
control
DMSO
cytochalasin D
**
**
G
DF-MDCK
Rupture Force (pN)
100
80
60
40
20
control
DMSO
cytochalasin D
**
**

**Supplementary figure 5** Plot of the median values with the 25th and 75th percentile added as error bars of rupture forces in cell-cell interactions between several cells under control, DMSO and cytochalasin D (5 µM) conditions. (A) NF-NF, (B) SF-SF, (C) DF-DF, (D) MDCK-MDCK, (E) NF-MDCK, (F) SF-MDCK and (G) DF-MDCK. The respective median values are also listed in Table 2. Statistical results are reported in Materials and Methods section.

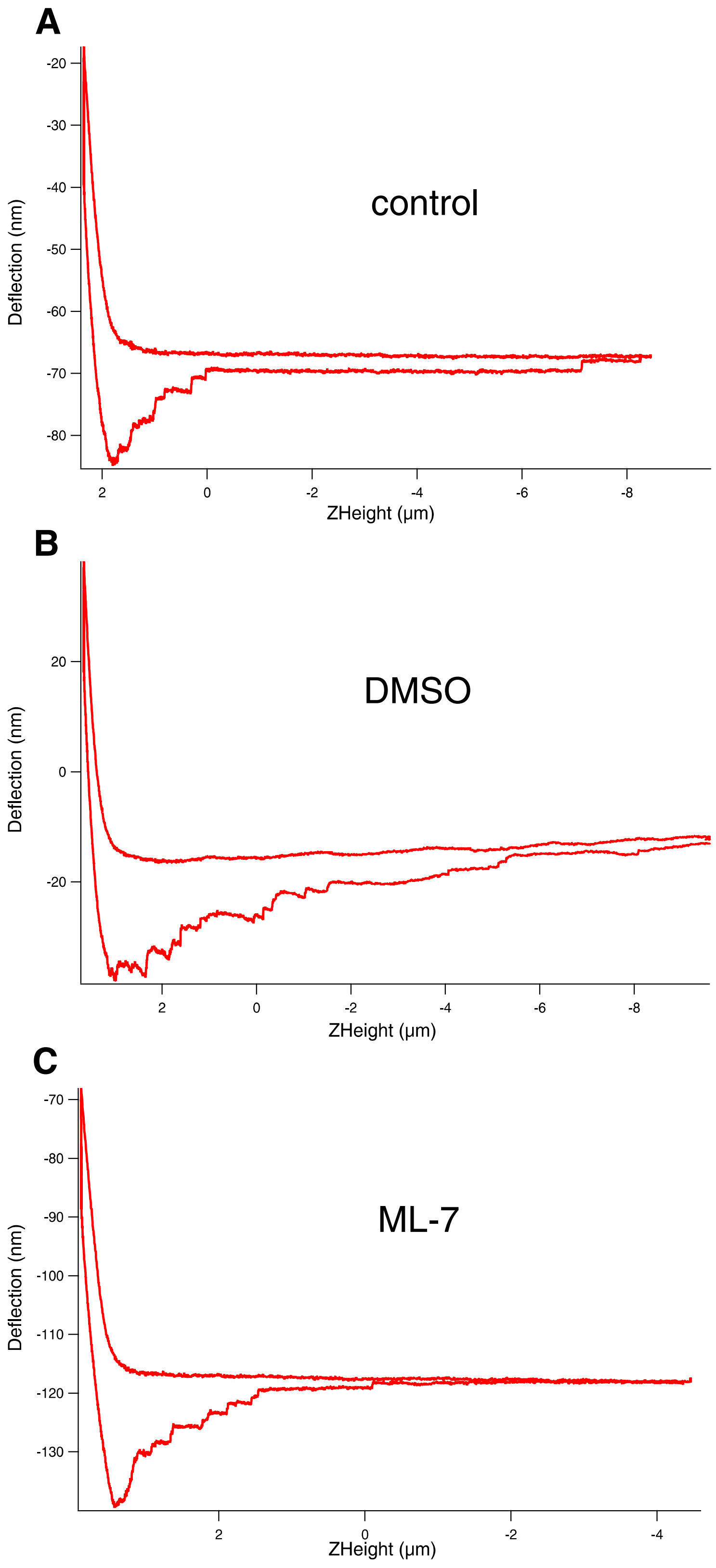

**Supplementary figure 6** Force curves measured during cell-cell interaction of normal fibroblasts (NF-NF) shows rupture events in all three experimental conditions: control in normal medium (A), medium plus DMSO (B), and medium with ML-7 (C).

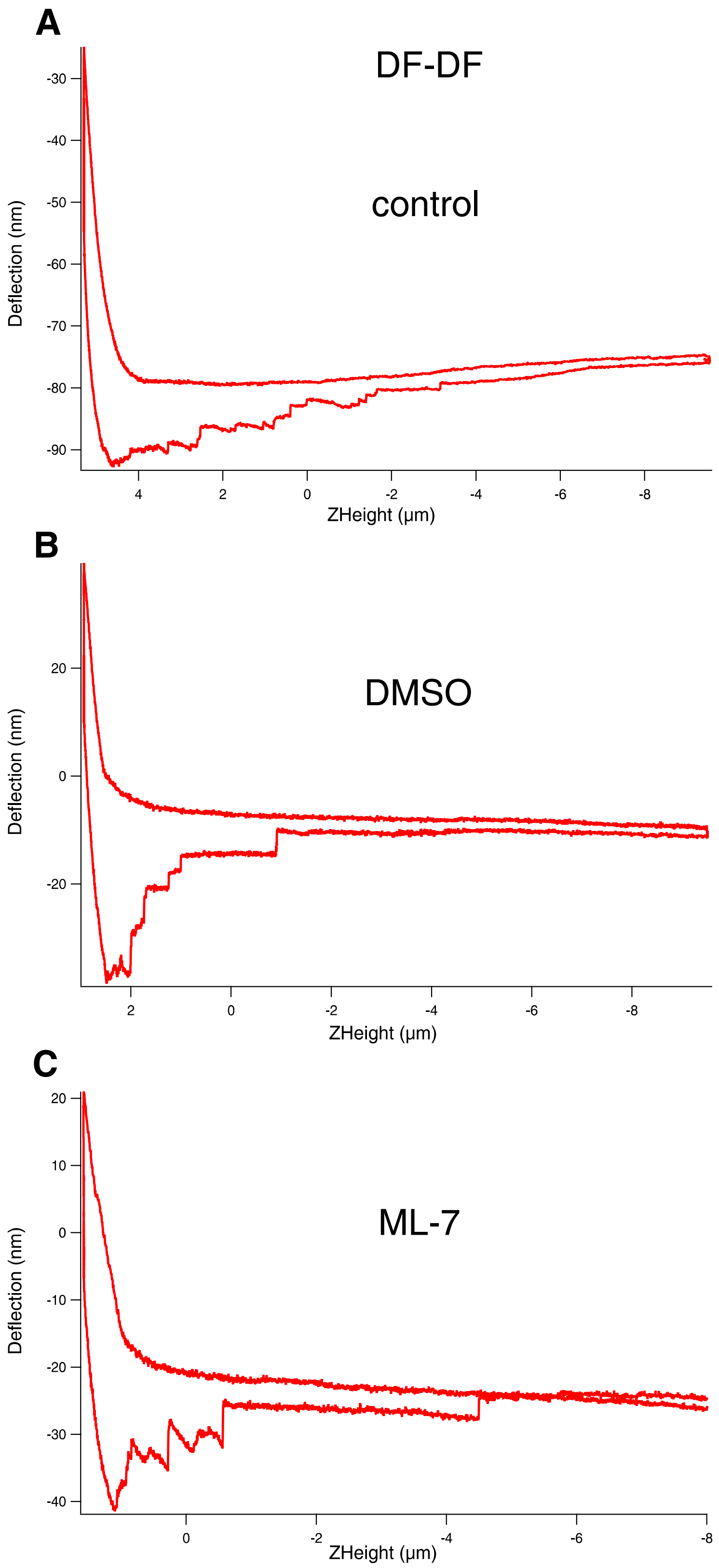

**Supplementary figure 7** Force curves measured during cell-cell interaction of Dupuytren fibroblasts (DF-DF) shows rupture events in all three experimental conditions: control in normal medium (A), medium plus DMSO (B), and medium with ML-7 (C).

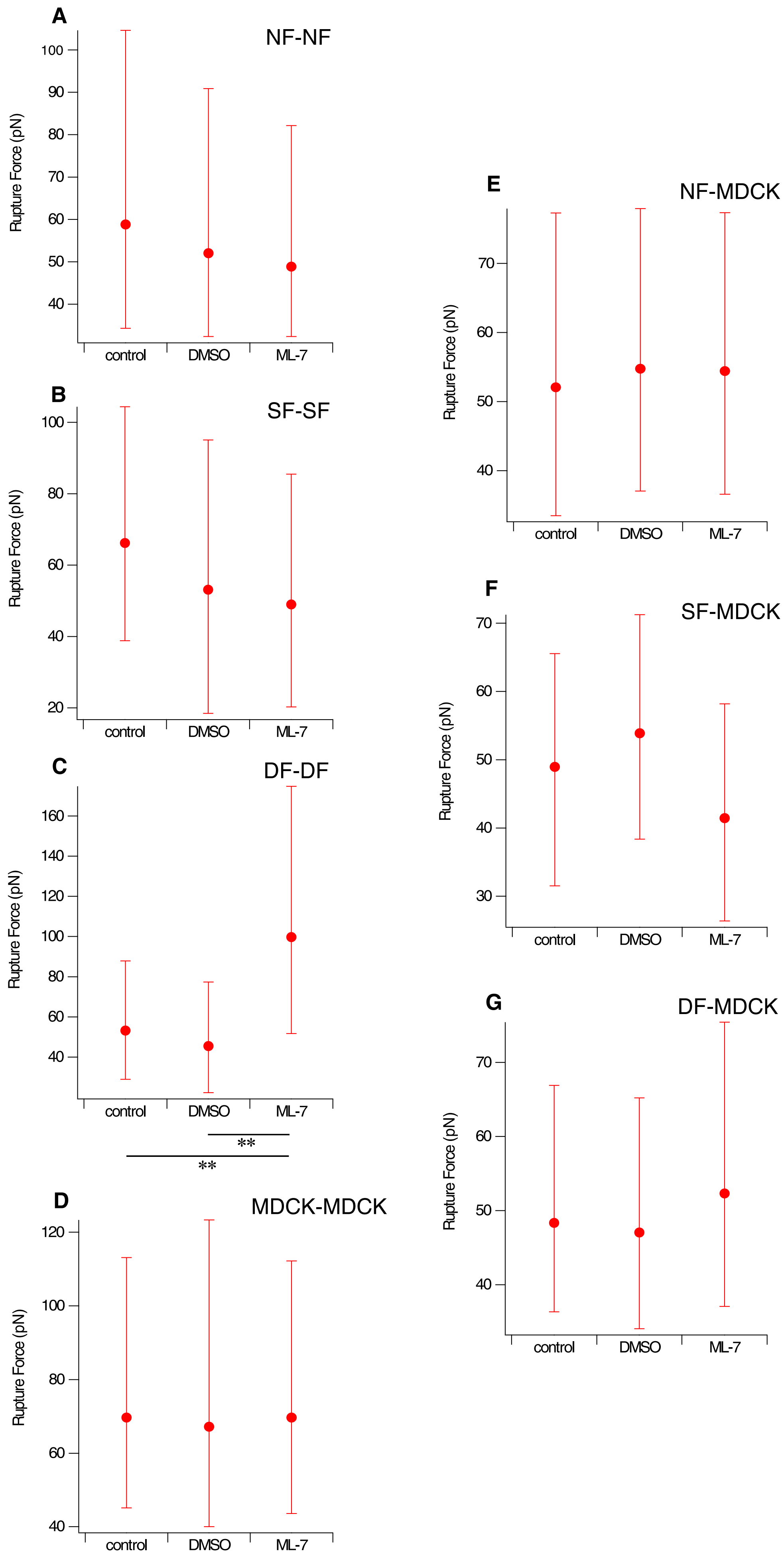
A
NF-NF
Rupture Force (pN)
100
90
80
70
60
50
40
control
DMSO
ML-7
B
SF-SF
Rupture Force (pN)
100
80
60
40
20
control
DMSO
ML-7
C
DF-DF
Rupture Force (pN)
160
140
120
100
80
60
40
control
DMSO
ML-7
**
**
D
MDCK-MDCK
Rupture Force (pN)
120
100
80
60
40
control
DMSO
ML-7
E
NF-MDCK
Rupture Force (pN)
70
60
50
40
control
DMSO
ML-7
F
SF-MDCK
Rupture Force (pN)
70
60
50
40
30
control
DMSO
ML-7
G
DF-MDCK
Rupture Force (pN)
70
60
50
40
control
DMSO
ML-7

**Supplementary figure 8** Plot of the median values with the 25th and 75th percentile added as error bars of rupture forces in cell-cell intaractions between several cells under control, DMSO and ML-7 (5 μM) conditions. (A) NF-NF, (B) SF-SF, (C) DF-DF, (D) MDCK-MDCK, (E) NF-MDCK, (F) SF-MDCK and (G) DF-MDCK. The respective median values are also listed in Table 3. Statistical results are reported in Materials and Methods section.
